# Adipocyte Kindlin-2 Coordinates an E-cadherin/β-catenin/IGFBP2 Endocrine Axis to Regulate Skin Regeneration and Diabetic Wound Healing

**DOI:** 10.64898/2026.11.01.763554

**Authors:** Zhen Ding, Guixing Ma, Fen Wang, Siyuan Cheng, Pengxuan Li, Yong Chen, Litong Chen, Milei Ouyang, Aimin Wu, Haobo Pan, Huiling Cao

## Abstract

Skin wound healing is a complex, multicellular process essential for maintaining tissue integrity and homeostasis. Despite advances in biomaterial and cell-based therapies, the mechanisms by which adipose tissue regulates skin regeneration remain incompletely understood. Here, we identify adipocyte Kindlin-2 as a critical regulator of cutaneous wound repair. Adipocyte-specific deletion of Kindlin-2 markedly accelerates wound closure in mice by enhancing fibroblast proliferation, angiogenesis, and suppressing inflammation. Transplantation of visceral white adipose tissue (vWAT) from Kindlin-2–deficient mice similarly promotes wound healing, indicating an endocrine contribution of vWAT to skin regeneration. Mechanistically, loss of Kindlin-2 promotes E-cadherin proteolysis, resulting in increased release of a 33-kDa C-terminal fragment (CTF2) that activates β-catenin signaling and induces transcriptional upregulation and secretion of insulin-like growth factor binding protein 2 (IGFBP2). Elevated circulating IGFBP2, in turn, stimulates integrin signaling in target cells to enhance wound repair. Pharmacological inhibition of β-catenin signaling abrogated the healing advantage conferred by Kindlin-2 deficiency. Conversely, local delivery of IGFBP2 via lentiviral significantly accelerates wound closure in both wild-type C57BL/6 and diabetic *ob/ob* mice. Collectively, these findings uncover a previously unrecognized adipose–skin endocrine axis governed by Kindlin-2 and IGFBP2 that orchestrates tissue regeneration. Targeting this Kindlin-2/E-cadherin/β-catenin/IGFBP2 pathway offers a promising therapeutic strategy for enhancing wound healing, particularly in chronic and diabetic wounds.

## Introduction

The skin serves as the body’s first line of defense, protecting against dehydration, mechanical trauma, ultraviolet radiation, microbial invasion, and thermal stress (1). Cutaneous wounds—particularly chronic ulcers and burns—pose a major threat to health and impose substantial socioeconomic burdens. In the United States, approximately 0.75% of the population suffers from pressure ulcers, and among more than 30 million individuals with diabetes, nearly one million develop diabetic foot ulcers each year (2). It is estimated that 6-7 million people will experience a foot ulcer during their lifetime (3), with annual healthcare costs associated with acute and chronic skin wounds exceeding $50 billion (4).

Skin repair is a highly orchestrated, dynamic process traditionally divided into three overlapping stages: inflammation, proliferation, and remodeling (5). Although the skin possesses remarkable regenerative capacity, complete restoration of its structure and function is often compromised under pathological conditions such as extensive burns, chronic wounds, aging, and diabetes (6–9). Current therapeutic strategies— including conventional dressings, bioactive scaffolds, cell-based therapies, and autologous grafts—have achieved limited success, largely due to heterogeneity in wound pathology and patient variability (1). Therefore, elucidating the regulatory mechanisms governing cutaneous repair and identifying novel therapeutic targets remain of critical importance.

White adipose tissue (WAT), composed of adipocytes, preadipocytes, immune cells, endothelial cells, and stromal cells, functions not only as an energy reservoir but also as the body’s largest endocrine organ (10). Through the secretion of diverse adipokines—including proteins, microRNAs, lipids, exosomes—WAT plays a fundamental role in systemic metabolic homeostasis (11–14). Recent studies have revealed that dermal white adipose tissue (dWAT) is an active participant in skin homeostasis and wound healing (15, 16). Dermal adipocytes dedifferentiate and migrate into the wound bed as progenitor-like cells (16, 17), while paracrine factors derived from these cells modulate keratinocyte proliferation, fibroblast activation, and immune responses to coordinate tissue regeneration (18). However, whether other adipose depots—particularly visceral white adipose tissue (vWAT)—also influence cutaneous wound healing through endocrine mechanisms remains largely unknown.

Kindlin-2, encoded by *Fermt2*, is a focal adhesion protein widely expressed in mesenchymal-derived tissues, where it regulates cell adhesion, migration, survival, and differentiation (19, 20). Previous studies, including our own, have demonstrated that Kindlin-2 is indispensable for the development and homeostasis of multiple organs, including bone, cartilage, liver, pancreas, heart, and kidney (21–29). We previously reported that adipocyte-specific Kindlin-2 deletion impairs WAT expansion by reducing adipocyte size (30), suggesting that adipocyte Kindlin-2 is a key regulator of metabolic homeostasis—a process intricately linked to skin repair and regeneration.

Insulin-like growth factor binding protein 2 (IGFBP2), a member of the IGFBP family, binds IGF-I and IGF-II to modulate their bioactivity and signaling activity (31, 32). Beyond its canonical IGF-dependent effects, IGFBP2 exerts IGF-independent actions on cell migration, proliferation, angiogenesis, and stemness (33–35), mediated in part through interactions with extracellular matrix proteins such as integrins (36–38). Although IGFBP2 contributes to skin development, its role in cutaneous wound healing remains unclear.

Here, we identify adipocyte Kindlin-2 as a critical regulator of skin wound repair. Adipocyte-specific deletion of Kindlin-2 markedly accelerates wound closure by promoting fibroblast proliferation, angiogenesis, and attenuating inflammation. Transplantation of vWAT from adipocyte-specific Kindlin-2 knockout mice similarly promotes wound healing, underscoring the endocrine contribution of vWAT to cutaneous regeneration. Mechanistically, Kindlin-2 loss augments E-cadherin proteolysis, leading to increased generation of a 33-kDa C-terminal intracellular fragment (CTF2) that activates β-catenin signaling and upregulates IGFBP2 expression and secretion. Circulating IGFBP2, in turn, facilitates wound repair by stimulating the IGFBP2-integrin signaling cascade. Notably, local administration of IGFBP2 significantly accelerates cutaneous healing in both wild-type C57BL/6 and diabetic *ob/ob* mice. Together, these findings uncover a previously unrecognized adipose–skin endocrine axis that governs tissue regeneration and identify IGFBP2 as a promising therapeutic target for chronic wound management.

## Results

### Adipocyte-specific Kindlin-2 deletion accelerates skin wound healing

To investigate the role of adipocyte Kindlin-2 in skin wound healing, we generated adipocyte-specific Kindlin-2 knockout mice (*Adi-Cre; Kindlin-2^fl/fl^*, hereafter referred to as KO), as we previously described (30). Cre-negative littermates (*Kindlin-2^fl/fl^*) served as controls (hereafter referred to as Con). Western blot analysis (WB) confirmed efficient and specific deletion of Kindlin-2 in adipose tissues (Fig. S1 A and B). As expected, Kindlin-2 deficiency markedly reduced fat mass, particularly in vWAT, without affecting overall body weight and other major organ weights (Fig. S1 C and D).

To assess the impact of adipocyte Kindlin-2 loss on skin wound repair, we employed a full-thickness excision model (8-mm diameter) and monitored wound closure every two days. KO mice displayed a significantly accelerated wound closure rate compared with control mice, achieving nearly complete re-epithelialization by day 8 (Fig. 1A and B). Histological analyses revealed a narrower wound gap and increased callus thickness in KO mice (Fig. 1C and D). Masson’s trichrome staining further demonstrated enhanced collagen deposition at the wound sites of KO mice (Fig. 1E and F).

**Figure 1.**
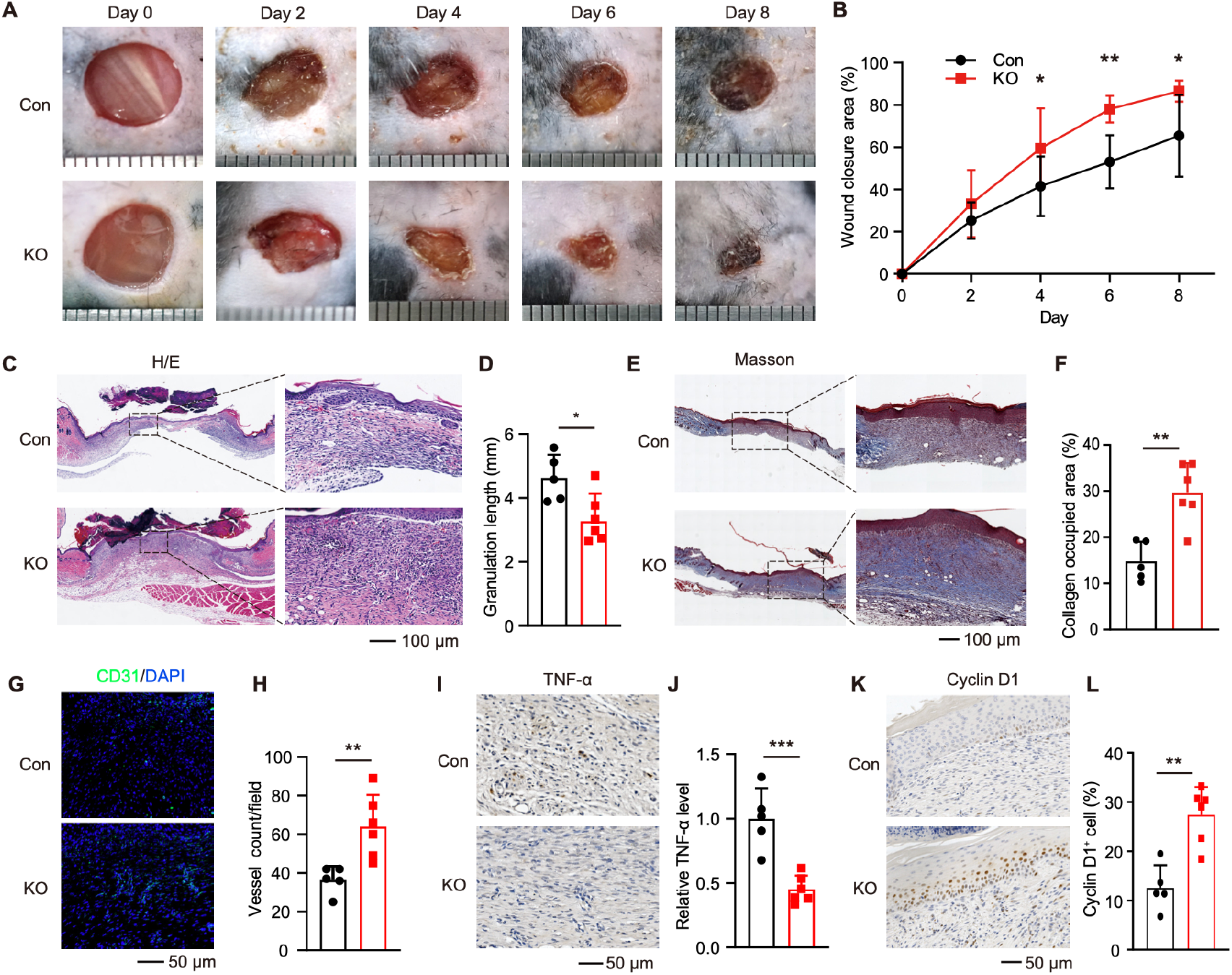
Adipocyte-specific Kindlin-2 deficiency markedly enhances skin wound healing. **A**. Representative images of full-thickness skin wounds in adipocyte-specific Kindlin-2 knockout (KO) and control (Con) mice at indicated time points (0, 2-, 4-, 6-, and 8-days post-surgery). Scale bars, 1 mm. **B**. Quantification of wound closure area in Con and KO mice. **C-F**. Histological analysis of skin tissue sections collected 9 days after surgery. Representative H/E staining (C) and quantification of granulation tissue length (D). Representative Masson’s trichrome staining (E) and quantifications of collagen deposition area (F). **G** and **H**. Immunofluorescence (IF) staining for CD31 in wound sections at day 8 post-surgery. Representative images (G) and quantifications of vessel density (average vessels per field for every mouse) (H). **I-L**. Immunohistochemical staining of wound sections at day 9 post-surgery showing (I) TNF-α and (K) Cyclin D1 expression, with corresponding quantifications of (J) relative TNF-α levels and (L) Cyclin D1–positive cell percentage. Data are presented as mean ± SD (n = 5 Con mice, n = 6 KO mice). Statistical significance was determined by unpaired two-tailed Student’s *t*-test (D, F, H, G, and L) or two-way ANOVA (F). \**P* < 0.05; \*\**P* < 0.01; \*\*\**P* < 0.001; \*\*\*\**P* < 0.0001.

Given the importance of neovascularization in tissue repair(39), we next assessed angiogenesis by immunofluorescence (IF) staining for CD31, a marker of vascular endothelial cells. Wounds from KO mice exhibited significantly increased angiogenesis (Fig. 1G and H). Immunohistochemical (IHC) staining for TNF-α indicated a reduced inflammatory response in KO wounds (Fig. 1I and J), while Cyclin D1 staining revealed elevated cell proliferation (Fig. 1K and L), consistent with enhanced tissue regeneration (40).

Collectively, these results demonstrate that adipocyte Kindlin-2 plays a critical inhibitory role in skin wound healing, and its loss promotes accelerated repair through enhanced angiogenesis, collagen deposition, and cellular proliferation.

### vWAT grafts from KO mice accelerate skin wound healing

To determine whether distal vWAT influences skin wound healing, we transplanted vWAT from 5-month-old KO or Con mice into 3-month-old wild-type (WT) C57BL/6 recipients. One month after transplantation, full-thickness excisional skin wounds were created, and wound closure was monitored every two days (Fig. 2A). IF staining for Kindlin-2 confirmed the survival and persistence of transplanted adipose tissue (Fig. 2B and C).

**Figure 2.**
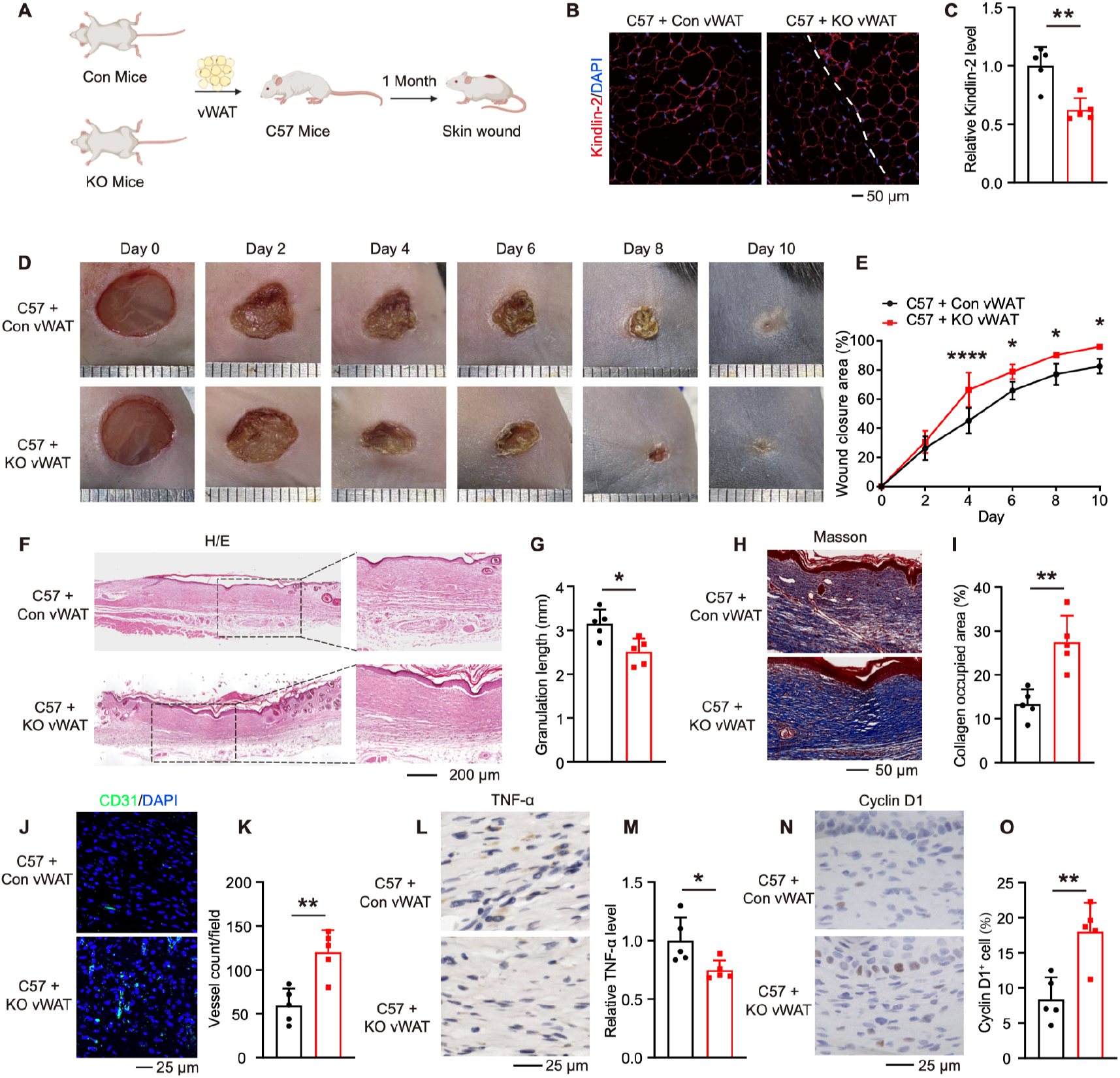
vWAT grafts from KO mice accelerates skin wound healing. **A**. Schematic illustration of the experimental design evaluating the effect of vWAT grafts on wound healing. **B** and **C**. Immunofluorescence (IF) staining of vWAT sections from wild-type C57BL/6 mice transplanted with Con vWAT (C57 + Con vWAT) or KO vWAT (C57 + KO vWAT). Representative images of Kindlin-2 staining (B) and quantification of relative Kindlin-2 levels (C). **D.** Representative images of full-thickness skin wounds in C57 + Con vWAT and C57 + KO vWAT mice at indicated time points (0, 2-, 4-, 6-, 8-and 10-days post-surgery). Scale bars, 1 mm. **E.** Quantification of wound closure area over time. **F-I**. Histological analysis of skin tissue sections collected 10 days post-surgery. Representative H/E staining (F) and quantification of granulation tissue length (G). Representative Masson’s trichrome staining (H) and quantification of collagen deposition area (I). **J** and **K.** IF staining for CD31 in wound sections at day 10 post-surgery. Representative images (J) and quantification of vessel density (average vessels per field for every mouse) (K). **L-O**. Immunohistochemical staining of wound sections at day 10 post-surgery showing (L) TNF-α and (N) Cyclin D1 expression, with corresponding quantifications of (M) relative TNF-α levels and (O) Cyclin D1–positive cell percentage. Data are presented as mean ± SD (n = 5 mice for per group). Statistical significance was determined by unpaired two-tailed Student’s *t*-test (C, G, I, K, M, and O) or two-way ANOVA (E). \**P* < 0.05; \*\**P* < 0.01; \*\*\**P* < 0.001; \*\*\*\**P* < 0.0001.

Notably, mice receiving vWAT from KO donors exhibited significantly accelerated wound closure compared with those receiving grafts from control mice (Fig. 2D and E). Histological analyses using H/E and Masson’s trichrome staining revealed enhanced wound callus formation and greater collagen deposition in wounds of the KO vWAT group (Fig. 2F-I). Moreover, IHC staining for CD31, TNF-α, and Cyclin D1 demonstrated increased neovascularization, reduced inflammation, and elevated cell proliferation at the wound sites of mice transplanted with KO vWAT (Fig. 2J-O). Notably, mice receiving vWAT grafts exhibited comparable body weights between groups (Fig. S2A).

Collectively, these findings indicate that vWAT from KO mice markedly promotes cutaneous wound repair, highlighting the role of adipose tissue-derived systemic signals in skin regeneration.

### IGFBP2 derived from vWAT promotes skin wound regeneration through activation of integrin signaling

To identify adipose-derived factors responsible for the enhanced wound repair observed in KO mice, we isolated mRNA from the vWAT of KO and control mice for transcriptomic analysis. The top 30 up-regulated genes in the KO group were identified, and cross-referencing these with secreted factors from the Human Protein Altas and SEPDB databases yielded 5 significantly upregulated candidates (Fig. 3A).

**Figure 3.**
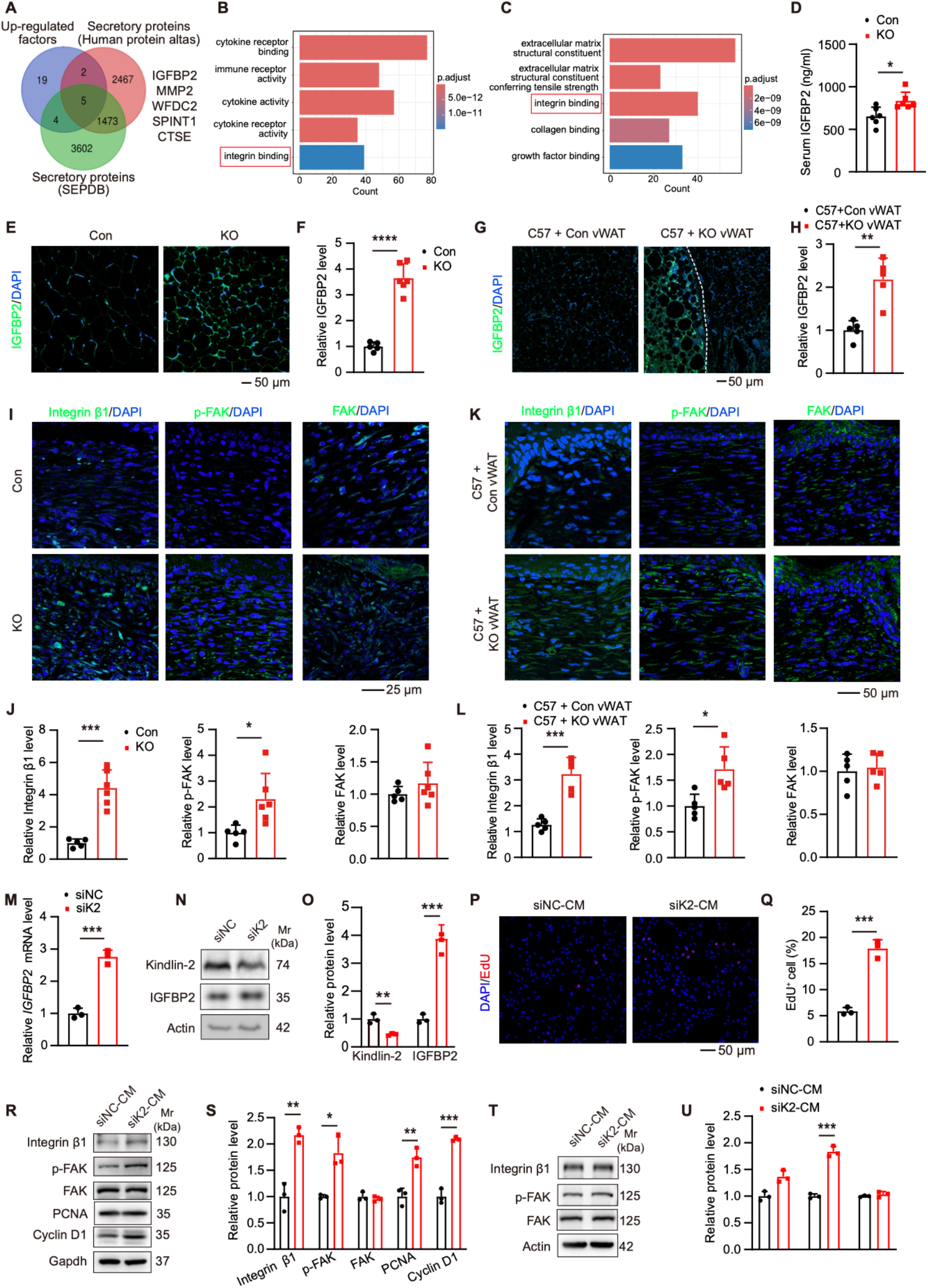
IGFBP2 derived from vWAT promotes skin wound regeneration. **A**. Venn diagram showing the overlap among the top 30 up-regulated factors in KO vWAT and the secretory protein databases Human Protein Atlas and SEPDB. **B** and **C**. KEGG pathway analysis of significantly altered pathways in skin tissues from (B) injured mice (GSE23006) and (C) injured humans (GSE199568). **D**. ELISA assay measuring serum IGFBP2 levels in Con and KO mice. **E** and **F**. IF staining of IGFBP2 in vWAT from Con and KO mice. Representative images (E) and quantification of IGFBP2 fluorescence intensity (F). **G** and **H**. IF staining of IGFBP2 in vWAT from C57BL/6 mice transplanted with Con vWAT (C57 + Con vWAT) or KO vWAT (C57 + KO vWAT). Representative images (G) and quantification of IGFBP2 levels (H). **I** and **J**. IF staining of healing skin tissue from C57BL/6 mice transplanted with Con or KO vWAT. Representative images showing Integrin β1, p-FAK, and total FAK (I); and corresponding quantification of fluorescence intensity (J). **K** and **L**. IF staining of healing skin tissue from C57 + Con vWAT and C57 + KO vWAT groups. Representative images showing Integrin β1, p-FAK, and FAK (K) and quantification of fluorescence intensity (L). **M-O**. Differentiated 3T3-L1 adipocytes transfected with Kindlin-2 siRNA (siK2) or negative control siRNA (siNC) were analyzed by (M) RT-qPCR for IGFBP2 mRNA expression and by (N–O) Western blotting (WB) for IGFBP2 protein levels. **P-S**. NIH3T3 cells were cultured for 48 h with 30% conditioned medium (CM) from siNC adipocytes (siNC-CM) or siK2 adipocytes (siK2-CM). Representative EdU staining images (P), quantification of proliferating cells (Q), WB analysis (R), and quantification of protein expression (S). **T** and **U**. HUVECs were cultured for 48 h with 30% CM from siNC or siK2 adipocytes. WB analysis (T) and quantification of protein expression (U). Data are presented as mean ± SD. For (D, F, J), n = 5 Con mice, n = 6 KO mice. For (H, L), n = 5 mice per group. For (M, O, Q, S, U), n = 3 biologically independent experiments. Statistical significance was determined by unpaired two-tailed Student’s *t*-test. \**P* < 0.05; \*\**P* < 0.01; \*\*\**P* < 0.001; \*\*\*\**P* < 0.0001.

To further explore signaling pathways relevant to skin regeneration, we re-analyzed publicly available RNA-seq datasets of mouse (GEO: GSE23006) (41, 42) and human (GEO: GSE199568) (43) skin wounds (Fig. 3B and C)). Integrin signaling emerged as one of the most significantly enriched pathways in both species (Fig. 3B and C), consistent with prior reports highlighting its essential role in skin repair and angiogenesis (44, 45). Among the five secreted candidates, insulin-like growth factor– binding protein 2 (IGFBP2)—a regulator of integrin activation (46)—was particularly notable.

ELISA assays revealed that plasma IGFBP2 levels were markedly elevated in KO mice compared to controls (Fig. 3D), and similarly increased in recipients transplanted with KO vWAT (Fig. S3A). Both IF staining and RT-qPCR confirmed significantly higher IGFBP2 expression in vWAT from KO mice at both the mRNA and protein levels (Fig. 3E and F and Fig. S3B). Consistently, KO-derived vWAT grafts retained higher IGFBP2 expression than control grafts or native vWAT in recipient mice (Fig. 3G and H).

Previous studies have shown that IGFBP2 exerts its biological functions by regulating Integrin α5β1 heterodimers and their downstream signaling pathways (47, 48). Moreover, Integrin β1 and its downstream effector FAK have been identified as key mediators of skin wound healing (44, 45). To determine whether integrin signaling is activated in response to IGFBP2 elevation, we examined Integrin β1 and FAK activation in skin wounds. IF staining revealed robust increases in Integrin β1 and phosphorylated FAK (p-FAK) in the wounds of KO mice relative to controls (Fig. 3I and J), while total FAK levels remained unchanged (Fig. 3I and J). Similarly, enhanced Integrin β1 and p-FAK expression were detected in wound tissues of mice receiving KO vWAT grafts (Fig. 3K and L). Collectively, these results indicate that Kindlin-2 deficiency in adipocytes enhances IGFBP2 production, leading to systemic IGFBP2 elevation and subsequent activation of integrin signaling and to promote tissue regeneration.

To further validate the role of adipocyte-derived IGFBP2, we knocked down Kindlin-2 in 3T3-L1 preadipocytes and collected conditioned medium (CM) after adipogenic differentiation. RT-qPCR and WB analyses confirmed that Kindlin-2 silencing significantly upregulated IGFBP2 expression (Fig. 3M-O). Treatment of NIH3T3 fibroblasts with CM from Kindlin-2–deficient adipocytes (siK2-CM) markedly increased Integrin β1 and p-FAK levels compared with control CM (siNC-CM) (Fig. 3R and S). Consistently, fibroblast proliferation was enhanced, as evidenced by elevated Cyclin D1 expression and increased EdU incorporation (Fig. 3P-S). Moreover, HUVECs treated with siK2-CM displayed augmented Integrin β1 and p-FAK expression, indicative of enhanced integrin signaling (Fig. 3T and U).

Together, these findings demonstrate that adipocyte Kindlin-2 deficiency promotes IGFBP2 secretion from vWAT, leading to systemic activation of integrin signaling, which stimulates fibroblast proliferation and angiogenesis—mechanisms that likely underlie the accelerated wound healing phenotype observed in KO mice.

### Adipocyte Kindlin-2 deficiency activates E-cadherin/β-catenin signaling to induce IGFBP2 expression

To elucidate the molecular mechanism by which Kindlin-2 regulates IGFBP2 expression, we examined the involvement of β-catenin, a known transcriptional regulator of IGFBP2 (49). IF staining revealed markedly increased β-catenin protein levels in the vWAT of KO mice compared with controls (Fig. 4A and B).

**Figure 4.**
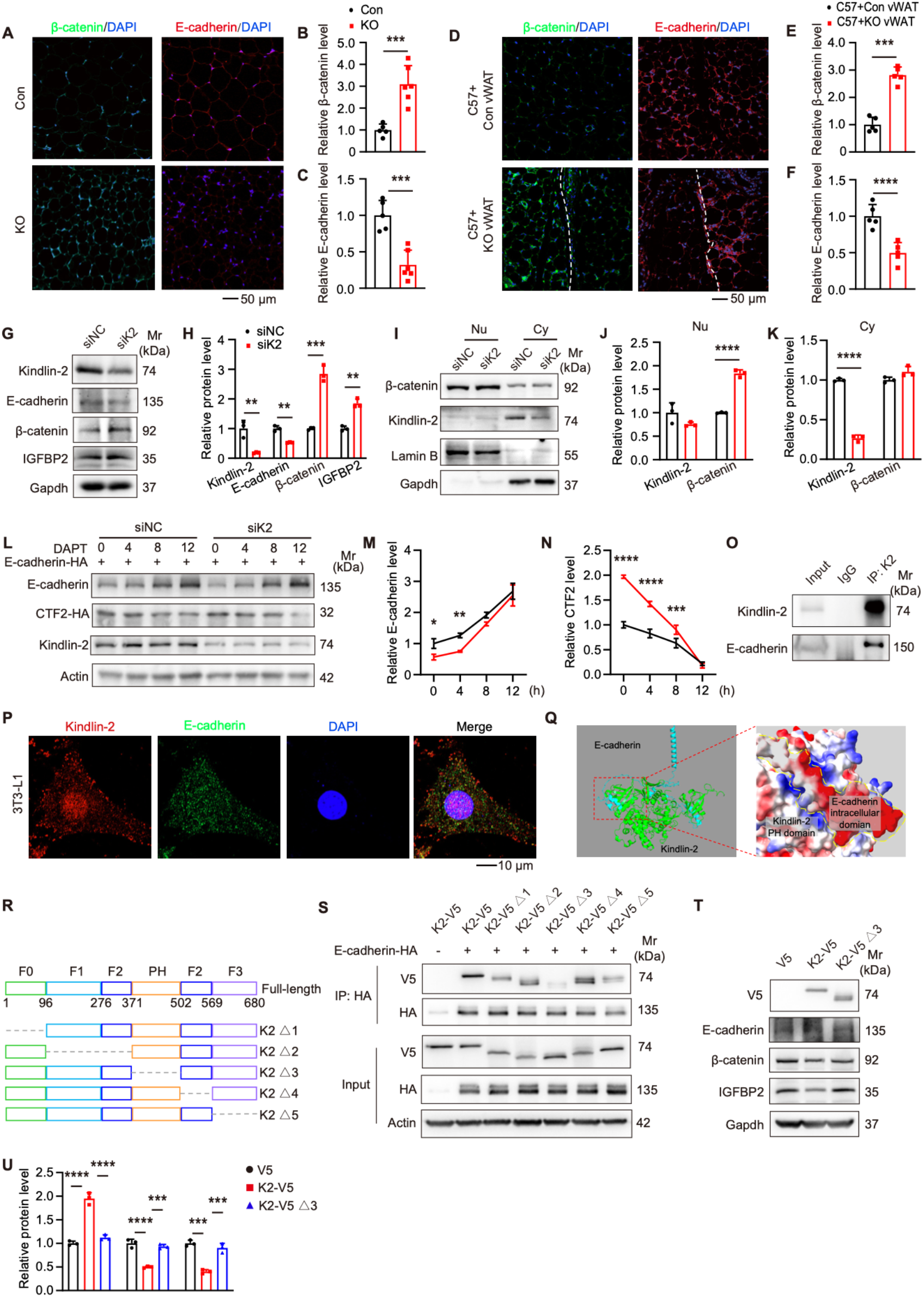
Kindlin-2-deficiency enhances E-cadherin/β-catenin signaling to promote IGFBP2 expression in adipocytes. **A-C**. IF staining of β-catenin and E-cadherin in vWAT from Con and KO mice. Representative images (A) and quantification of fluorescence intensity (B, C). **D-F**. IF staining of β-catenin and E-cadherin in vWAT from C57BL/6 mice transplanted with Con vWAT (C57 + Con vWAT) or KO vWAT (C57 + KO vWAT). Representative images (D) and quantification of fluorescence intensity (E, F). **G** and **H**. Differentiated 3T3-L1 adipocytes transfected with Kindlin-2 siRNA (siK2) or control RNA (siNC) were analyzed by WB analysis (G) and corresponding quantification (H). **I-K**. Differentiated 3T3-L1 adipocytes transfected with siK2 or siNC were harvested after 72 h, and nuclear (Nu) and cytoplasmic (Cy) proteins were separately extracted for WB analysis. (I) Representative WB images and quantification of (J) nuclear and (K) cytoplasmic protein levels. **L-N**. Adipocytes transfected with siK2 or siNC and overexpressing E-cadherin-HA were treated with DAPT and collected at different time points for WB analysis. Representative WB images (L) and quantification of relative E-cadherin levels (normalized to siNC, 0 h) (J) and relative CTF2 levels (normalized to siNC, 0 h) (K). **O**. Co-IP analysis of Kindlon-2 and E-cadherin interaction in 3T3-L1 preadipocytes. **P**. IF staining showing colocalization of Kindlin-2 (red) and E-cadherin (green) in 3T3-L1. **Q**. Predicted structural model of the Kindlin-2 and E-cadherin interaction. **R**. Schematic representation of Kindlin-2 truncation mutants. K2 Δ1 lacks the FERM0 (F0) domain, K2 Δ2 lacks the F1 and F2-1 domains, K2 Δ3 lacks the PH domain, K2 Δ4 lacks the F2-2 domain, and K2 Δ5 lacks the F3 domain. **S**. Co-IP analysis of Kindlin-2 truncation mutants and E-cadherin interaction in 3T3-L1. **T** and **U**. Differentiated 3T3-L1 adipocytes transfected with control plasmid (V5), Kindlin-2 overexpression plasmid (K2-V5), or PH-domain-truncated Kindlin-2 plasmid (K2-V5 Δ3) were analyzed by WB analysis (T) and quantification of protein expression (U). Data are presented as mean ± SD. For (B, C), n = 5 Con mice, n = 6 KO mice. For (E, F), n = 5 mice per group. For (H, J, K, M, N, U), n = 3 biologically independent experiments. *P* values were determined by unpaired two-tailed Student’s *t*-test (B, C, E, F, H, J, K), two-way ANOVA (M, N) or one-way ANOVA (U). \**P* < 0.05; \*\**P* < 0.01; \*\*\**P* < 0.001; \*\*\*\**P* < 0.0001.

E-cadherin, a key component of adherens junctions, binds β-catenin through its cytoplasmic domain and thereby modulates β-catenin signaling (50). Proteolytic cleavage of full-length E-cadherin releases a C-terminal fragment (CTF2), which promotes β-catenin nuclear translocation and transcriptional activation (51). IF staining demonstrated significantly reduced E-cadherin and increased β-catenin levels in the vWAT of KO mice (Fig. 4A and C). Consistently, vWAT grafts from KO donors exhibited elevated β-catenin and reduced E-cadherin expression compared with control grafts (Fig. 4D-F). RT-qPCR analysis revealed markedly increased *E-cadherin* mRNA expression, whereas *β-catenin* mRNA levels remained unchanged in the vWAT of KO mice relative to controls (Fig. S4A). These findings indicate that Kindlin-2 deletion in adipocytes upregulates E-cadherin transcription but decreases its protein abundance, likely through enhanced proteolytic cleavage, while β-catenin is regulated at the protein rather than mRNA level. Our observations are consistent with previous studies showing that β-catenin can enhance E-cadherin transcription (52, 53).

In vitro, differentiated 3T3-L1 adipocytes with Kindlin-2 knockdown displayed decreased E-cadherin and increased β-catenin and IGFBP2 protein levels, as confirmed by WB analysis (Fig. 4G and H). Furthermore, Kindlin-2 knockdown significantly enhanced β-catenin nuclear translocation (Fig. 4I-K).

Collectively, these results indicate that loss of Kindlin-2 in adipocytes promotes β-catenin activation by enhancing E-cadherin cleavage, leading to upregulation of IGFBP2 expression.

### Kindlin-2 interacts with E-cadherin and prevents its γ-secretase–mediated cleavage

To elucidate how Kindlin-2 regulates E-cadherin protein levels and downstream signaling, we examined whether Kindlin-2 influences E-cadherin stability or proteolytic processing, given that Kindlin-2 is neither an enzyme nor a transcription factor. We overexpressed C-terminally HA-tagged E-cadherin in differentiated 3T3-L1 adipocytes and treated the cells with the proteasome inhibitor MG132. WB analysis revealed that Kindlin-2 knockdown reduced full-length E-cadherin levels while concomitantly increasing CTF2, regardless of MG132 treatment (Fig. S4B and C). These results suggest that both full-length E-cadherin and CTF2 are degraded via the proteasome, but loss of Kindlin-2 primarily enhances CTF2 generation rather than altering its degradation.

Cycloheximide (CHX) chase assays further demonstrated that Kindlin-2 knockdown did not alter the half-life of either full-length E-cadherin or its C-terminal fragment (CTF2), but markedly increased CTF2 production (Fig. S4D-F). Notably, treatment with the γ-secretase inhibitor DAPT completely abolished the accumulation of CTF2 induced by Kindlin-2 knockdown (Fig. 4L-N), indicating that Kindlin-2 inhibits γ-secretase–mediated E-cadherin cleavage.

We next investigated whether Kindlin-2 interacts with E-cadherin to exert this regulatory effect. Co-immunoprecipitation (Co-IP) confirmed an endogenous interaction between Kindlin-2 and E-cadherin (Fig. 4O), while IF staining revealed their co-localization in differentiated 3T3-L1 adipocytes (Fig. 4P). To predict the interaction interface, we employed AlphaFold-Multimer modelling using full-length Kindlin-2 and the cytoplasmic domain of E-cadherin. The predicted structure indicated a potential binding interface between the cytoplasmic domain of E-cadherin and the PH domain of Kindlin-2, stabilized by electrostatic interactions (Fig. 4Q).

To validate this prediction experimentally, we performed Co-IP assays using a series of Kindlin-2 truncation mutants (Fig. 4R). Only the mutant lacking the PH domain (K2-V5 Δ3) failed to co-precipitate with E-cadherin (Fig. 4S), confirming that the PH domain of Kindlin-2 is essential for its interaction with E-cadherin. Furthermore, overexpression of full-length Kindlin-2 in 3T3-L1 adipocytes decreased E-cadherin levels while increasing β-catenin and IGFBP2 expression, whereas overexpression of the PH domain–deleted mutant had no effect (Fig. 4T-U).

Collectively, these findings demonstrate that the PH domain of Kindlin-2 mediates its interaction with E-cadherin, thereby suppressing γ-secretase–dependent E-cadherin cleavage and regulating β-catenin signaling and IGFBP2 expression in adipocytes.

### β-catenin inhibition reverses the enhanced wound healing phenotype caused by adipocyte Kindlin-2 deficiency

To investigate whether the promotion of IGFBP2 expression and wound healing by Kindlin-2 deficiency depends on β-catenin signaling, we employed the specificβ-catenin inhibitor ICG001. WB analysis showed that ICG001 significantly suppressed β-catenin and its downstream target IGFBP2 expression in adipocytes, and effectively reversed the upregulation of both proteins induced by Kindlin-2 deficiency (Fig. 5A and B).

**Figure 5.**
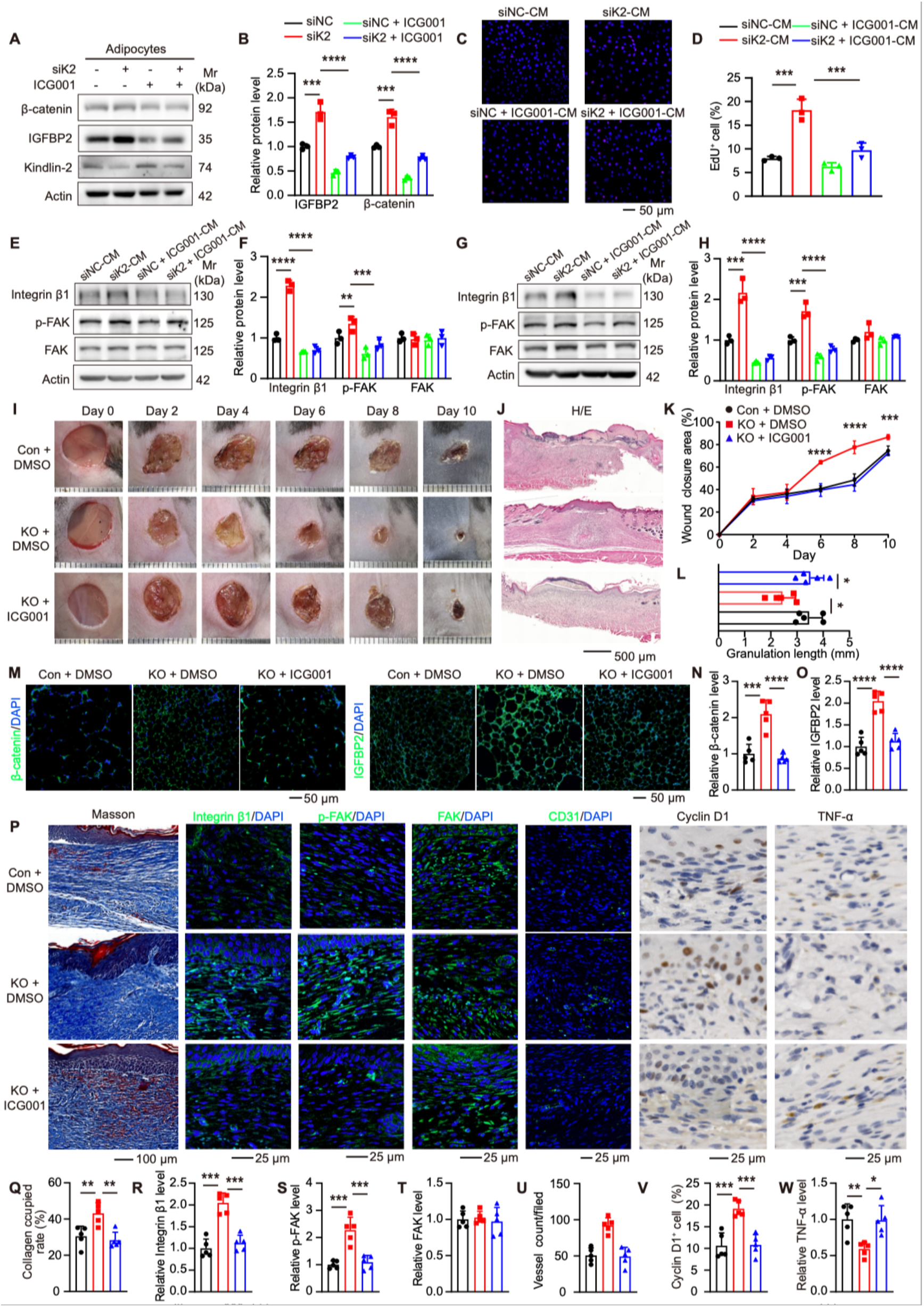
β-catenin inhibitor ICG-001 reverses the promotive effect of Kindlin-2 deficiency on skin wound healing. **A** and **B**. Differentiated 3T3-L1 adipocytes were transfected with siK2 or siNC and treated with or without ICG001 for WB analysis (A) and quantification (B), and conditioned medium (CM) was collected after 48 h. **C-F**. NIH3T3 cells were cultured for 48 h in medium containing 30% CM. Representative images of EdU staining (C) and quantification of proliferating cells (D). WB analysis of NIH3T3 cells (E) and corresponding quantification (F). **G** and **H**. HUVECs were cultured for 48 h in medium containing 30% CM. WB analysis (G) and quantification (H). **I**. Representative images of full-thickness skin wounds from control mice treated with DMSO (Con + DMSO), KO mice treated with DMSO (KO + DMSO), and KO mice treated with ICG-001 (KO + ICG-001) at the indicated time points (0, 2-, 4-, 6-, 8-, and 10-days post-surgery). Scale bars, 1 mm. **J**. Representative H/E-stained sections of wound tissue at day 10 post-surgery. **K**. Quantification of wound closure area. **L**. Quantification of granulation tissue length. **M-O**. IF staining of vWAT from Con + DMSO, KO + DMSO, and KO + ICG-001 mice. (M) Representative images of β-catenin and IGFBP2 and quantification of (N) β-catenin and (O) IGFBP2 fluorescence intensity. **P-W**. Histological and immunostaining analyses of skin wound sections from Con + DMSO, KO + DMSO, and KO + ICG-001 mice. Representative images of Masson’s trichrome staining, IF and IHC (P). Quantifications of collagen deposition area (Q), Integrin β1 (R), p-FAK (S), total FAK (T), CD31 positive vessel density (average vessels per field for every mouse) (U), Cyclin D1 positive cell percentage (V), and TNF-α levels (W). Data are presented as mean ± SD. For (B, D, F, H), n = 3 biologically independent experiments. For (K, L, N, O, Q-W), n = 5 mice per group. *P* values were determined by two-way ANOVA (B, D, F, H) or one-way ANOVA (K, L, N, O, Q-W,). \**P* < 0.05; \*\**P* < 0.01; \*\*\**P* < 0.001; \*\*\*\**P* < 0.0001.

Conditional medium (CM) collected from mature adipocytes was next applied to NIH3T3 fibroblasts. CM from control adipocytes treated with ICG001 (siNC + ICG001-CM) significantly reduced Integrin β1 and p-FAK levels as well as cell proliferation. Likewise, CM derived from Kindlin-2-deficient adipocytes treated with ICG001 (siK2 + ICG001-CM) abolished the siK2-CM activation of integrin signaling, as confirmed by WB analysis and EdU staining (Fig. 5C-F). Consistently, in HUVECs, the elevated Integrin β1 and p-FAK levels observed with siK2-CM were reversed by siK2 + ICG001-CM treatment (Fig. 5G and H). These results indicate that Kindlin-2 regulates IGFBP2 expression through β-catenin signaling *in vitro*.

To determine whether adipocyte Kindlin-2 regulates IGFBP2 secretion through β-catenin signaling in vivo, 5-month-old female KO mice were treated with the β-catenin inhibitor ICG001 for three weeks, followed by the induction of full-thickness excisional skin wounds. ICG001 treatment did not affect body weight or organ mass in KO mice (Fig. S5A and B). As expected, KO mice exhibited accelerated wound closure between days 6 and 10 post-injury compared with control mice; however, ICG001 treatment completely reserved this enhanced healing phenotype (Fig. 5I-L). Ten days after injury, adipose and wound tissues were collected for analysis. ICG001 treatment significantly reduced β-catenin and IGFBP2 expression in vWAT of KO mice (Fig. 5M-O). Histological analysis further showed that ICG-001attenuated wound callus thickness and collagen deposition (Fig. 5P and Q). IF and IHC staining demonstrated that ICG001 treatment diminished Integrin β1 and p-FAK expression, reduced CD31-positive vessel density, suppressed Cyclin D1 expression, and restored TNF-α levels in KO mice (Fig. 5P and R-W).

Together, these findings demonstrate that β-catenin inhibition reserves the accelerated wound healing induced by adipocyte Kindlin-2 deficiency, indicating that Kindlin-2 deficiency promotes IGFBP2 expression and enhances skin wound repair primarily through activation of β-catenin signaling.

### IGFBP2 promotes skin wound healing in WT C57BL/6 mice

Although IGFBP2 is a potent regulator of integrin signaling—an essential pathway for skin repair (44)—its direct role in wound healing remains unclear. To investigate this, we constructed a lentiviral vector overexpressing IGFBP2 (Lv-IGFBP2) and assessed its effects on fibroblast activity and skin regeneration.

In NIH3T3 fibroblasts, Lv-IGFBP2 infection markedly increased IGFBP2 protein abundance, elevated Integrin β1 levels, and enhanced FAK phosphorylation compared with control virus (Lv-NC) (Fig. S6A and B). Correspondingly, IGFBP2 overexpression significantly promoted fibroblast proliferation, as evidenced by increased Cyclin D1 and PCNA expression and enhanced EdU incorporation (Fig. S6A-D). Similarly, in HUVECs, IGFBP2 overexpression also led to elevated Integrin β1 and p-FAK levels (Fig. S6E and F).

To evaluate the therapeutic potential of IGFBP2 *in vivo*, full-thickness excisional wounds were generated on the dorsal skin of 5-month-old WT C57BL/6 mice. Lv-IGFBP2 was locally delivered within an alginate gel matrix at the wound site. Mice treated with Lv-IGFBP2 exhibited markedly accelerated wound closure between day 4 and 8 post-injury compared with Lv-NC–treated controls (Fig. 6A and B). By day 8, histological analysis revealed reduced wound width and increased granulation tissue thickness in Lv-IGFBP2–treated wounds (Fig. 6C and D). IF and IHC staining confirmed higher expression of IGFBP2, Integrin β1, and p-FAK in Lv-IGFBP2– treated wounds, accompanied by increased CD31-positive vessel density, elevated Cyclin D1 expression, and reduced TNF-α staining (Fig. 6E-L).

**Figure 6.**
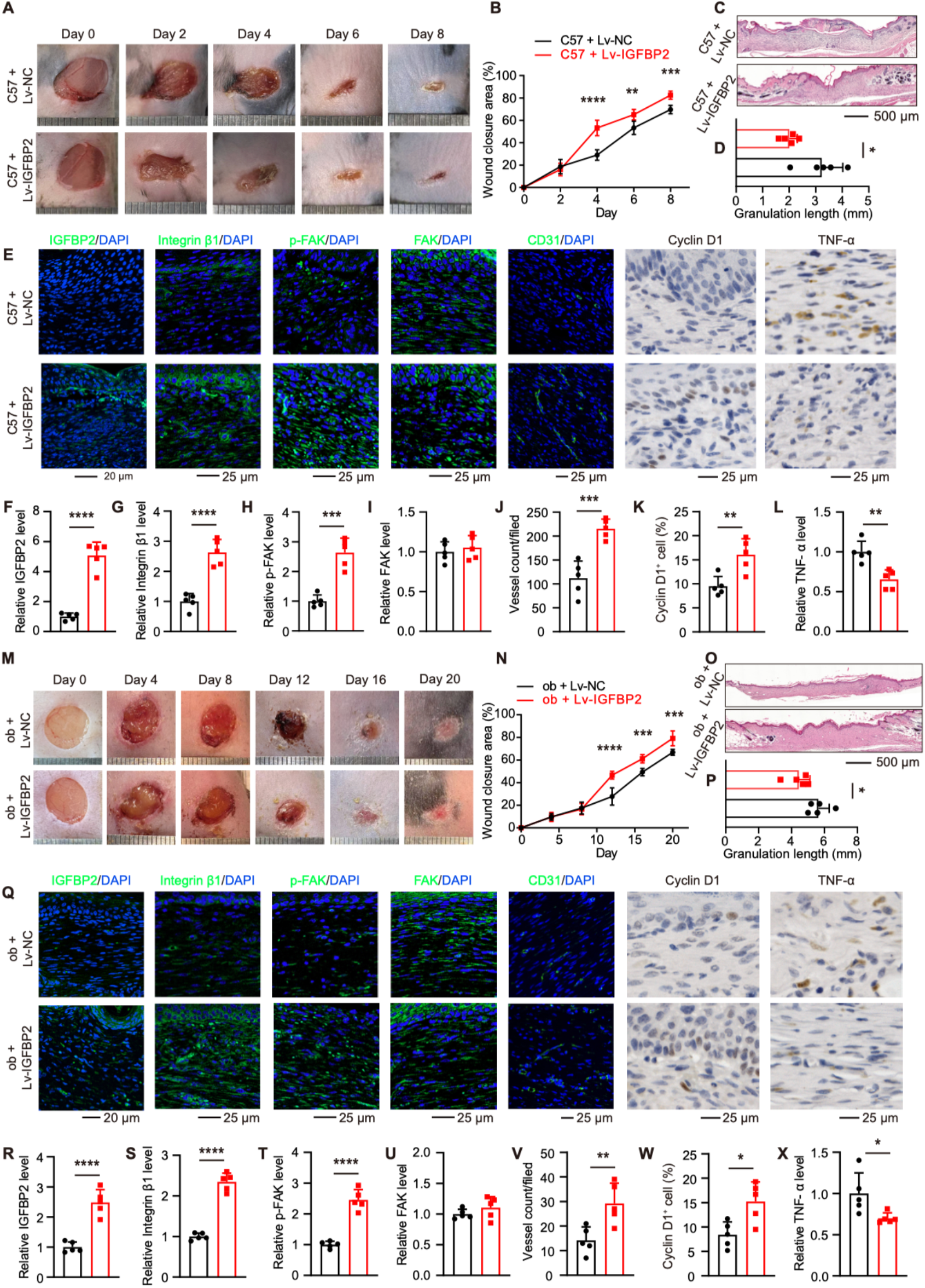
IGFBP2 markedly enhances skin wound healing in WT C57BL/6 mice and diabetic *ob/ob* mice. **A**. Representative images of full-thickness skin wounds in C57BL/6 mice treated with control lentivirus (C57 + Lv-NC) or IGFBP2-overexpressing lentivirus (C57 + Lv-IGFBP2) at the indicated time points (0, 2-, 4-, 6-, and 8-days post-surgery). Scale bars, 1 mm. **B**. Quantification of wound closure area. **C** and **D**. Representative H/E staining (C) and quantification of granulation tissue length (D) from C57 + Lv-NC and C57 + Lv-IGFBP2 mice at day 9 post-surgery. **E-L**. IF and IHC analyses of wound sections from C57 + Lv-NC and C57 + Lv-IGFBP2 mice. Quantification of (F) IGFBP2, (G) Integrin β1, (H) phosphorylated FAK (p-FAK), (I) total FAK, (J) CD31 positive vessel density, (K) Cyclin D1 positive cell percentage, and (L) TNF-α expression levels. **M**. Representative images of full-thickness skin wounds in diabetic *ob/ob* mice treated with control lentivirus (ob + Lv-NC), or IGFBP2 overexpressing lentivirus (ob + Lv-IGFBP2) at the indicated time points (0, 4-, 8-, 12-, 16-, 20-days after surgery). Scale bars, 1 mm. **N**. Quantification of wound closure area. **O** and **P**. Representative H/E staining (O) and quantification of granulation tissue length (P) from ob + Lv-NC and ob + Lv-IGFBP2 mice at day 20 post-surgery. **Q-X**. IF and IHC analyses of wound sections from ob + Lv-NC and ob + Lv-IGFBP2 mice. Quantification of (R) IGFBP2, (S) Integrin β1, (T) p-FAK, (U) total FAK, (V) CD31 positive vessel density average vessels per field for every mouse, (W) Cyclin D1 positive cell percentage, and (X) TNF-α expression levels. Data are presented as mean ± SD, n = 5 mice per group. *P* values were determined by unpaired two-tailed Student’s *t*-test or two-way ANOVA (B and N). \**P* < 0.05; \*\**P* < 0.01; \*\*\**P* < 0.001; \*\*\*\**P* < 0.0001.

Together, these findings demonstrate that IGFBP2 overexpression recapitulates the enhanced wound healing phenotype observed in KO mice, suggesting that local delivery of IGFBP2 represents a promising strategy to promote skin regeneration.

### IGFBP2 promotes wound healing in diabetic *ob/ob* mice

Chronic nonhealing skin wounds are a major complication of diabetes, for which effective treatments remain limited (54). *ob/ob* mice, which lack leptin, are widely used as a model of type 2 diabetes and display markedly delayed wound healing (55). IF staining revealed that Kindlin-2 expression in vWAT was significantly elevated in *ob/ob* mice compared with wild-type C57BL/6 controls (Fig. S6G and H). Notably, E-cadherin was drastically unregulated, whereas β-catenin and IGFBP2 were significantly downregulated in *ob/ob* mice relative to controls (Fig. S1G-K). These findings are consistent with our core observations and suggest that dysregulation of the Kindlin-2/E-cadherin/β-catenin/IGFBP2 axis may contribute to impaired wound healing under diabetic conditions.

To assess the therapeutic potential of IGFBP2 under diabetic conditions, full-thickness dorsal skin wounds were created in 5-month-old *ob/ob* mice, followed by local application of Lv-IGFBP2 embedded in an alginate gel matrix. Compared with control lentivirus (Lv-NC), Lv-IGFBP2 treatment significantly accelerated wound closure throughout the healing period (Fig. 6M and N). Histological analyses revealed thicker granulation tissue and enhanced collagen deposition in Lv-IGFBP2–treated wounds, as shown by H/E and Masson’s trichrome staining (Fig. 6O and P). Moreover, IHC and IF staining demonstrated increased CD31 positive vessel density, elevated IGFBP2, Integrin β1, p-FAK, and Cyclin D1 expression, and reduced TNF-α levels in the Lv-IGFBP2 treated group (Fig. 6Q-X).

Collectively, these results indicate that local IGFBP2 overexpression promotes wound healing in diabetic *ob/ob* mice by enhancing fibroblast proliferation and angiogenesis while suppressing inflammation, highlighting IGFBP2 delivery as a promising therapeutic strategy for diabetic wound repair.

## Discussion

In this study, we identify adipocyte Kindlin-2 as a pivotal regulator of skin wound healing and establish the E-cadherin/β-catenin/IGFBP2 axis as a key signaling pathway linking adipose tissue to skin regeneration. Our data reveal that adipocyte-specific loss of Kindlin-2 markedly accelerates wound closure by enhancing fibroblast proliferation, angiogenesis, and attenuating local inflammation. Mechanistically, Kindlin-2 directly interacts with E-cadherin in adipocytes and suppresses the proteolytic release of the E-cadherin C-terminal fragment (CTF2), thereby restraining β-catenin activation and its downstream effector IGFBP2 (49, 56, 57). Kindlin-2 deficiency disrupts this repression, resulting in elevated β-catenin activity and IGFBP2 secretion.

Although previous studies reported that Kindlin-2 promotes β-catenin stabilization and transcriptional activity in tumor cells (58) or in pancreatic β-cells(24), our findings demonstrate that in mature adipocytes, Kindlin-2 acts as a negative regulator of β-catenin signaling. The dual function of Kindlin-2 may be caused by different cell types and different microenvironment. This microenvironment-dependent function, which is common in biomolecules, for example IL-10, considered as an anti-inflammatory cytokine(59), can promote immune response via inducing B cell proliferation, survival and producing antibodies during some autoimmunity(60, 61). Prior studies showed that systemic haploinsufficiency of Kindlin-2 impairs skin wound healing by locally affecting angiogenesis (62), which is opposite to our current findings. This tissue-specific divergence highlights the functional versatility of Kindlin-2 and underscores its context-dependent roles across different cell types. Together with our previous findings that Kindlin-2 deficiency in adipocytes alters white adipose tissue (WAT) expansion (30), the current study further broadens the functional spectrum of adipocyte Kindlin-2, positioning it as a systemic coordinator of tissue regeneration and metabolic homeostasis.

Adipose tissue, particularly WAT, has emerged as an active regulator of skin repair. Dermal WAT (dWAT) supplies progenitors that transdifferentiate into myofibroblasts, while subcutaneous WAT (sWAT) contributes to granulation tissue formation and inflammatory responses (16, 63). Here, we extend this concept by demonstrating that visceral WAT (vWAT) also exerts a potent paracrine influence on skin wound healing through the secretion of IGFBP2. Transplantation of vWAT from adipocyte-specific Kindlin-2 knockout mice enhanced wound closure, emphasizing the significance of adipose-derived IGFBP2 as a systemic mediator connecting the WAT–skin axis.

IGFBP2, a multifunctional member of the IGFBP family, regulates adipogenesis, angiogenesis, and metabolism (37, 64). Our study uncovers a previously unrecognized role of IGFBP2 as a potent pro-healing factor that activates integrin signaling—an essential pathway for skin regeneration (44)—and markedly promotes wound repair. Importantly, IGFBP2 overexpression via lentiviral delivery accelerated wound closure not only in normal mice but also in diabetic *ob/ob* mice, a clinically relevant model of chronic, non-healing wounds. These findings highlight IGFBP2 as both a mechanistic link between adipose tissue and skin regeneration and a promising therapeutic candidate for treating impaired wound healing in metabolic disorders.

Despite these advances, we acknowledge that several limitations warrant further investigation. First, although our data demonstrate that IGFBP2 promotes wound healing through integrin activation, the precise downstream effectors and cellular targets within the wound microenvironment remain to be elucidated. Future studies using lineage-specific IGFBP2 receptor knockouts will help clarify these mechanisms. Second, the current study relies on lentiviral overexpression of IGFBP2. Evaluating the efficacy, safety, and delivery efficiency of recombinant IGFBP2 protein or small-molecule activators in preclinical models will be crucial for clinical translation. Collectively, this study establishes adipocyte Kindlin-2 as a central brake on the E-cadherin/β-catenin/IGFBP2 axis and reveals a novel adipose–skin regulatory mechanism controlling tissue repair. By delineating how adipocyte-derived IGFBP2 promotes fibroblast proliferation, angiogenesis, and inflammation resolution, our findings provide a conceptual advance in understanding the endocrine role of adipose tissue in tissue regeneration. Moreover, targeted activation of IGFBP2—either through gene therapy or recombinant protein delivery—holds strong translational potential for the treatment of chronic wounds, particularly in diabetic patients.

## Materials and Methods

### Animal Study

All animals used in this study were on a C57BL/6 genetic background and had been backcrossed for more than 10 generations. To achieve adipocyte-specific deletion of Kindlin-2, *Kindlin-2^fl/fl^* mice were crossed with *Adipoq-Cre* transgenic mice to generate *Adipoq-Cre; Kindlin-2^fl/fl^*(KO) mice, as previously described (30). Littermate *Cre*-negative *Kindlin-2^fl/fl^* mice were used as controls. All animals were housed under standard laboratory conditions (22–24 °C, 12 h light/12 h dark cycle) with ad libitum access to food and water. All animal procedures were approved by the Institutional Animal Care and Use Committee of Southern University of Science and Technology (SUSTech).

For the full-thickness skin wound model, 5-month-old KO and control mice or wild-type C57BL/6 mice were anesthetized with isoflurane, anddorsal hair was removed by shaving. Symmetrical full-thickness excisional wounds (8 mm in diameter) were created on the back using a sterile biopsy punch (65). Following surgery, each mouse was housed individually, and digital images of the wounds were captured on days 0, 2, 4, 6, 8, and 10 post-injury to monitor wound closure.

For adipose tissue transplantation, visceral white adipose tissue (vWAT) was collected from 5-month-old female control or KO donor mice. Approximately 2 g of freshly harvested vWAT was transplanted into the corresponding vWAT depot of 5-month-old female C57BL/6 recipient mice. One month after transplantation, full-thickness skin wounds were generated as described above.

For β-catenin inhibition, 5-month-old female KO mice received daily intraperitoneal injections of ICG-001 (S2662, Selleck) at a dose of 150 mg/kg body weight for three weeks. Control mice received vehicle (DMSO) injections (66). Full-thick skin wounds were then induced, and wound healing was assessed every two days thereafter.

For lentivirus treatment, skin wounds were treated immediately after surgery with lentivirus (Lv-NC or Lv-IGFBP2) at 2 × 10^9^ PFU/mL. After a 2-min incubation, wounds were covered with 1% alginate gel, and 150 mM CaCl₂ was applied to facilitate gel solidification (67). Each mouse was housed individually, and wound healing was monitored photographically every two days until closure.

### Cell Culture and Conditioned Medium Preparation

The 3T3-L1 preadipocyte cell line was maintained in Dulbecco’s modified Eagle’s medium (DMEM) supplemented with 10% fetal bovine serum (FBS; P30-3302, PAN Biotech), 100 U/mL penicillin, and 100 µg/mL streptomycin at 37 °C in a humidified incubator containing 5% CO₂.

For adipogenic differentiation, cells were cultured to confluence and then induced with adipogenic differentiation medium (PD-006, Procell) for 7 days. The conditional medium (CM) was collected at the end of differentiation, centrifuged to remove cell debris, and stored at −80 °C until use.

The NIH3T3 fibroblast cell line was maintained under the same culture conditions (DMEM supplemented with 10% FBS, 100 U/mL penicillin, and 100 µg/mL streptomycin) at 37 °C in 5% CO₂. For conditional medium treatment, 2×10^5^ NIH3T3 cells were seeded per well in 6-well plates and allowed to attach for 12 h. The medium was then replaced with fresh culture medium containing 30% differentiated 3T3-L1– derived conditioned medium, and cells were cultured for an additional 48 h before further analysis.

### Lentivirus Packaging and Preparation

Lentivirus production was performed as previously described (67). HEK293T cells were maintained in Dulbecco’s modified Eagle’s medium (DMEM) supplemented with 10% fetal bovine serum (FBS), 100 U/mL penicillin, and 100 µg/mL streptomycin at 37 °C in a humidified 5% CO₂ incubator. The target plasmid (Lv-IGFBP2 or Lv-NC) together with the packaging plasmids psPAX2 and pMD2.G were co-transfected into HEK293T cells using Lipofectamine 3000 reagent (L3000015, Invitrogen) according to the manufacturer’s instructions. The culture medium containing lentiviral particles was collected at 48 and 72 h post-transfection, followed by clarification and ultracentrifugation for viral concentration. The final lentiviral titer was approximately 2 × 10^9^ PFU/mL.

### Histological Analysis

Skin and adipose tissues were fixed in 4% paraformaldehyde for 24-48 h and embedded in paraffin. Paraffin blocks were sectioned at a thickness of 5 µm. Sections were deparaffinized, rehydrated, and subjected to hematoxylin and eosin (H/E) or Masson’s trichrome staining. For H/E staining, sections were stained sequentially with hematoxylin and eosin (BA4025, Baso), cleared in xylene, and mounted with neutral resin. For Masson’s trichrome staining, a commercial staining kit (G1340, Solarbio) was used according to the manufacturer’s instructions. Images were acquired using a Leica Aperio VERSA scanner and analyzed with ImageJ software. For quantification of Masson’s trichrome staining, the average across 6 fields from each mouse were analyzed, with 5 mice or 6 mice per group.

### Immunohistochemistry (IHC) and Immunofluorescence (IF)

Skin and adipose tissue were fixed in 4% paraformaldehyde for 24-48 h and embedded in paraffin. Paraffin blocks were sectioned at 5 µm thickness, deparaffinized, and subjected to antigen retrieval in citrate buffer.

For IHC staining, skin sections were incubated with a primary antibody against Cyclin D1 (Abclonal, A19038; 1:200), TNF-α (Abclonal, A11535; 1:200) at 4 ℃ overnight, followed by detection using the EnVision+System-HRP (DAB) kit (Dako North America Inc, K401111-2) according to the manufacturer’s instructions. Images were captured using a Lecia Aperio VERSA scanner and analyzed with ImageJ software. For quantification of IHC staining, the average across 6 fields from each mouse were analyzed, with 5 mice or 6 mice per group.

For IF staining, sections were incubated with primary antibodies against CD31 (Servicebio, GB153151; 1:100), Integrin β1(Abclonal, A21234; 1:100), phospho-FAK (Abclonal, A9032; 1:100), FAK (Abclonal, A11195; 1:100), Kindlin-2 (Millipore, MAB2617; 1:100), E-cadherin (Abclonal, A20798; 1:100), β-catenin (Abclonal, A19657; 1:100), and IGFBP2 (Abclonal, A9018; 1:200)) at 4 ℃ overnight, followed by incubation with fluorophore-conjugated secondary antibodies for 1 h at room temperature. Fluorescent images were acquired using a Zeiss LSM980 confocal microscope and analyzed with ImageJ software. For quantification of IHC staining, the average across 6 fields from each mouse were analyzed, with 5 mice or 6 mice per group.

### Western Blot Analysis

Cells or tissue were lysed in RIPA buffer (Sigma, R0278), and protein concertation was determined using a BCA protein assay kit (Cwbio, CW0014). Equal amounts of protein (20 µg per lane) were resolved by SDS–PAGE and transferred onto PVDF membranes. Membranse were blocked and then incubated with primary antibodies (Actin (ZSJQ, TA-09; 1:3000), Gapdh (ZSJQ, TA-08; 1:3000), Integrin β1(Abclonal, A21234; 1:1000), phospho-FAK (Abclonal, A9032; 1:1000), FAK (Abclonal, A11195; 1:1000), Kindlin-2 (Millipore, MAB2617; 1:1000), E-cadherin (Abclonal, A20798; 1:1000), β-catenin (Abclonal, A19657; 1:1000), IGFBP2 (Abclonal, A9018; 1:2000), Cyclin D1 (Abclonal, A19038; 1:1000), PCNA (Servicebio, GB14139; 1:1000), V5 (CST, 13202; 1:2000)) at 4 °C overnight, followed by incubation with HRP-conjugated secondary antibodies (anti-Mouse (ZSJQ, ZB-2305; 1:5000) and anti-Rabbit (ZSJQ, ZB-2301; 1:5000)) for 1 h at room temperature. Protein bands were visualized using an enhanced chemiluminescence substrate and quantified using ImageJ software.

### RNA Isolation and Quantitative Real-Time PCR (RT-qPCR)

Total RNA was extracted using an RNA isolation kit (Sangon, QI-10), and RNA concertation and purity were determined using a Nanodrop 3000 spectrophotometer. One microgram of total RNA was reverse transcribed into cDNA using a reverse transcription kit (Beyotime, D7190). RT-qPCR was performed using SYBR Green Master Mix (Beyotime, D7260) according to the manufacturer’s instructions. Relative mRNA expression levels were calculated using the ΔCt-method. All primers were synthesized by Sangon, and the primer sequences are listed in Table. S1.

### Quantification and Statistical Analysis

Mice were randomly assigned to experimental groups. IF and IHC analyses were performed in a double-blinded manner. Statistical analyses were conducted using GraphPad Prism 9 (Version 9.5.0). Data are presented as mean ± standard deviation (s.d.). Statistical significance was determined using two-tailed Student’s *t*-test, one-way ANOVA, or two-way ANOVA, as appropriate. A *p* value < 0.05 was considered statistically significant.

## Data, Materials, and Software Availability

All study data are included in the article and/or supporting information.

## Acknowledgments

We acknowledge the assistance of the Core Research Facilities at Southern University of Science and Technology. This work was supported in part by the National Key R&D Program of China (2024YFA0919200); the National Natural Science Foundation of China (82561160166, 824B2074, 825B2074, 82350710800, 82372476); the Shenzhen Medical Research Fund (B2504003); and the Guangdong Provincial Science and Technology Innovation Council (2017B030301018).

## Author Contributions

Z.D., G.M., H.P, and H.C. conceived and designed the study; Z.D., F.W., S.C., Y.C., P.L., L.C., M.O., A.W., and G.M. conducted the study, collected and analyzed the data; Z.D., G.M. and H.C. interpreted the data. Z.D., G.M. and H.C. drafted the manuscript. H.P, G.M. and H.C. are responsible for the integrity of the data analysis.

## Supplementary Tables

**Table S1.**
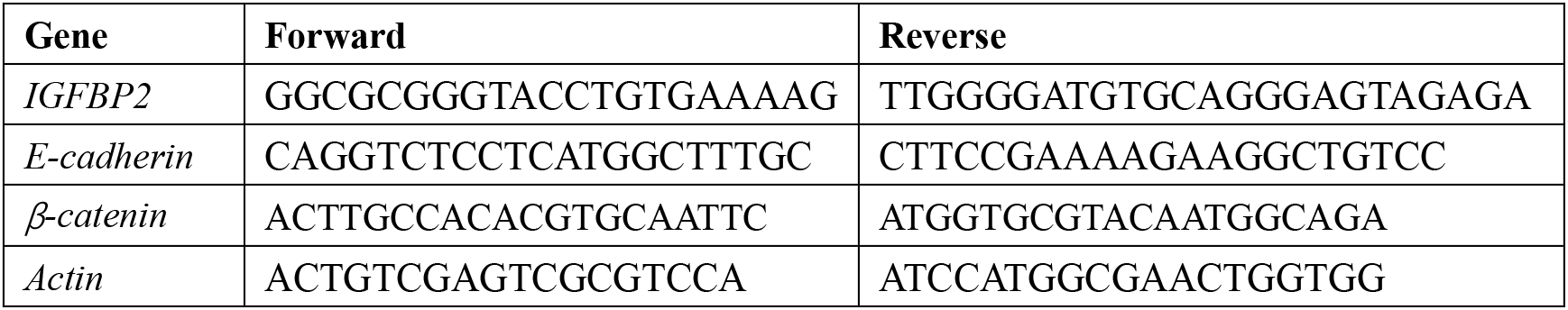
Primer information used in this study.

## Supplementary Figures

**Fig S1.**
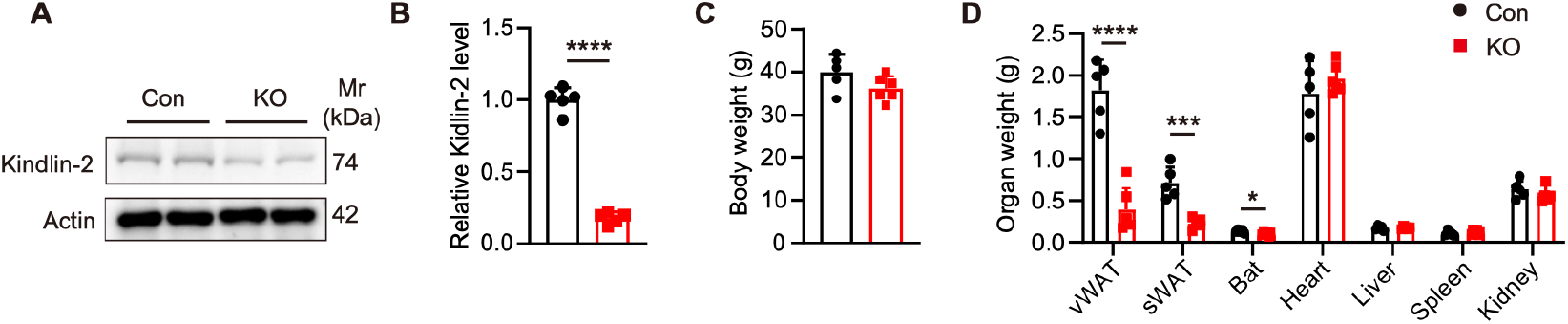
Adipocyte-specific Kindlin-2 deficiency reduces adipose tissue mass. **A** and **B**. Protein extracts from vWAT of adipocyte-specific Kindlin-2 knockout (KO) and control (Con) mice were analyzed by Western blotting (WB) to assess Kindlin-2 protein levels (A), with quantitative analysis shown in (B). **C**. Body weight of 5-month-old adipocyte-specific Kindlin-2 knockout (KO) and control mice. **D**. Organ weights of 5-month-old control an d KO mice. Data are presented as mean ± SD (n = 5 mice for control, n = 6 mice for KO). *P* values were determined using unpaired two-tailed Student’s *t*-test. \**P* < 0.05; \*\**P* < 0.01; \*\*\**P* < 0.001; \*\*\*\**P* < 0.0001.

**Fig S2.**
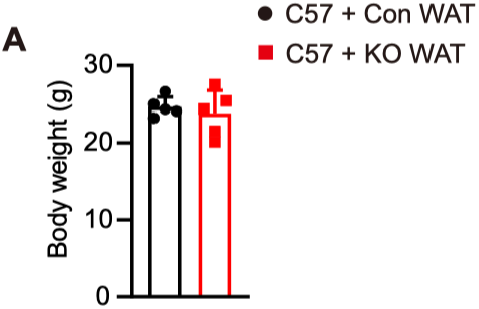
vWAT grafting does not affect body weight. **A.** Body weight of 3-month-old C57BL/6 mice transplanted with vWAT from control donors (C57 + Con vWAT) or KO donors (C57 + KO vWAT). Data are presented as mean ± SD (n = 5 mice per group). *P* values were determined by unpaired two-tailed Student’s *t*-test. \**P* < 0.05; \*\**P* < 0.01; \*\*\**P* < 0.001; \*\*\*\**P* < 0.0001.

**Fig S3.**
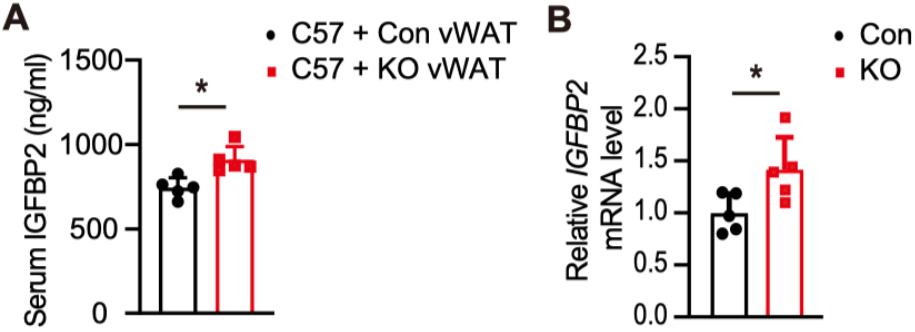
Adipocyte-specific Kindlin-2 deficiency increases IGFBP2 expression. **A.** ELISA quantification of serum IGFBP2 levels in C57BL/6 mice transplanted with control vWAT (C57 + WT vWAT) or KO vWAT (C57 + KO vWAT). **B.** RT-qPCR analysis for *IGFBP2* mRNA expression in vWAT from control and KO mice. Data are presented as mean ± SD (n = 5 mice per group). *P* values were determined using unpaired two-tailed Student’s *t*-test. \**P* < 0.05; \*\**P* < 0.01; \*\*\**P* < 0.001; \*\*\*\**P* < 0.0001.

**Fig S4.**
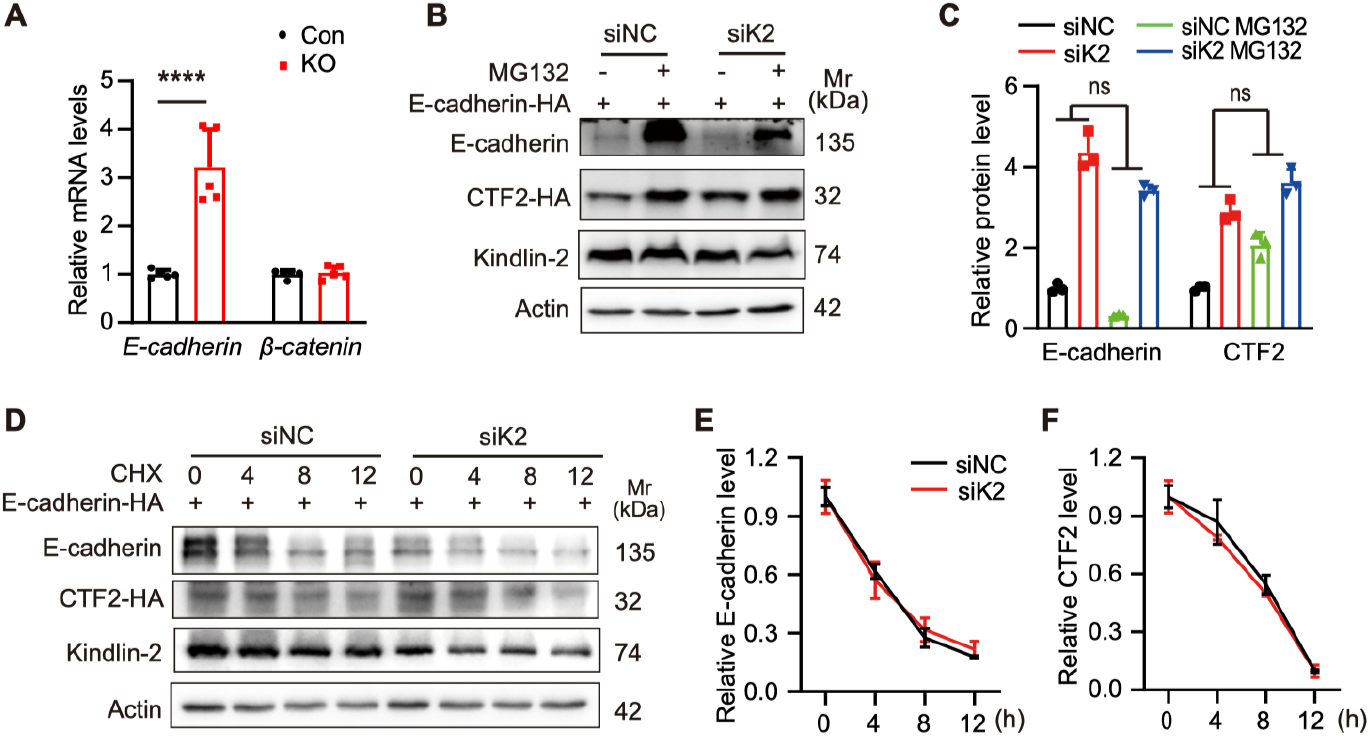
Kindlin-2 deficiency does not affect E-cadherin degradation. **A.** RT-qPCR analysis of vWAT from Con and KO mice showing mRNA levels of E-cadherin and β-catenin. **B** and **C.** Adipocytes transfected with siK2 or siNC were treated with MG132 or DMSO, and cell lysates were collected for WB analysis (B). Quantification of relative E-cadherin levels normalized to the siNC + DMSO group is shown in (C). **D-F**. Adipocytes transfected with siK2 or siNC and overexpressing E-cadherin-HA were treated with MG132 and harvested at the indicated time points for WB analysis (D). Quantifications of relative E-cadherin levels (E) and relative CTF2 levels were normalized to the 0 h level within each group (F). Data are presented as mean ± SD (n = 3 biologically independent experiments). *P* values were determined by two-way ANOVA (B) or one-way ANOVA (D and E). \**P* < 0.05; \*\**P* < 0.01; \*\*\**P* < 0.001; \*\*\*\**P* < 0.0001. ns, not significant.

**Fig. S5.**
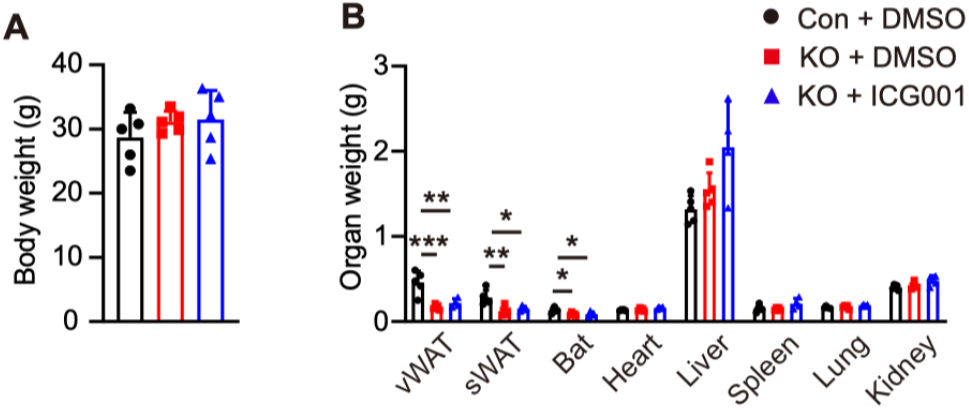
ICG001 treatment does not affect body weight and organ weights in mice. **A.** Body weight of 5-month-old control mice treated with DMSO (Con + DMSO), KO mice treated with DMSO (KO + DMSO), and KO mice treat with ICG001 (KO + ICG001). **B.** Organ weights of Con + DMSO, KO + DMSO, and KO + ICG001 mice. Data are presented as mean ± SD (n = 5 mice per group). *P* values were determined by one-way ANOVA. \**P* < 0.05; \*\**P* < 0.01; \*\*\**P* < 0.001; \*\*\*\**P* < 0.0001.

**Fig S6.**
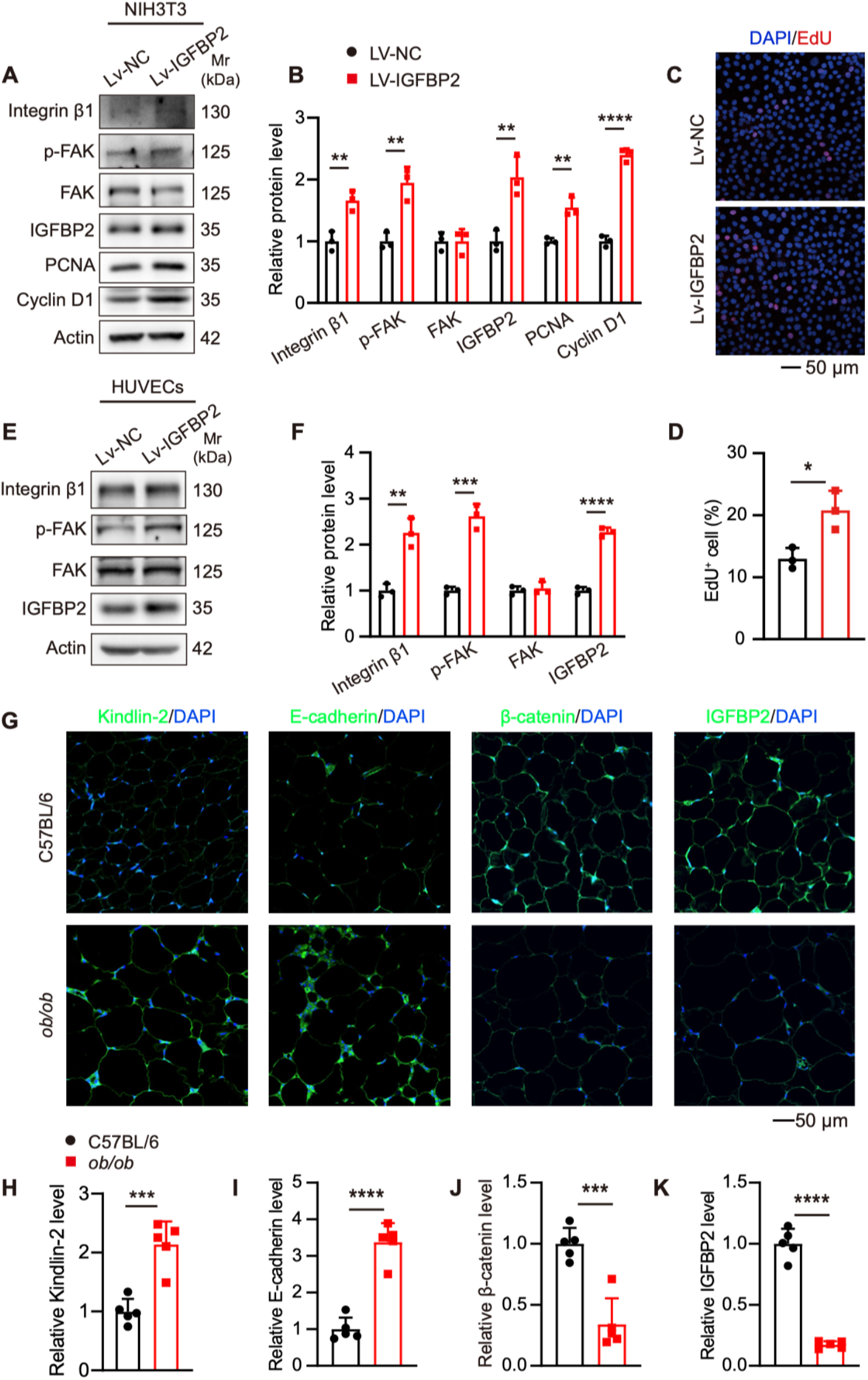
IGFBP2 overexpression activates integrin signaling and promotes cell proliferation. **A** and **B.** NIH3T3 cells were infected with control lentivirus (Lv-NC), or IGFBP2-overexpressing lentivirus (Lv-IGFBP2) and subjected to WB analysis (A) and quantification (B). **C** and **D**. NIH3T3 cells infected with Lv-NC or Lv-IGFBP2 were analyzed by EdU staining. Representative EdU images (C) and quantification of EdU-positive cells (D) are shown. **E** and **F**. HUVECs were infected with Lv-NC, or Lv-IGFBP2 and subjected to WB analysis (E) and quantification (F). **G-K**. IF staining of vWAT sections from 5-month-old C57BL/6 and *ob/ob* mice. Representative images showing Kindlin-2, E-cadherin, β-catenin and IGFBP2 expression are presented in (G), with quantification of Kindlin-2 (H), E-cadherin (I), β-catenin (J), and IGFBP2 (K). Data are presented as mean ± SD. For (B, D, F), n = 3 biologically independent experiments. For (H, I, J, K), n = 5 mice per group. *P* values were determined by unpaired two-tailed Student’s *t*-test. \**P* < 0.05; \*\**P* < 0.01; \*\*\**P* < 0.001; \*\*\*\**P* < 0.0001.

